# Bayesian MEG time courses with fMRI priors

**DOI:** 10.1101/2021.02.28.433293

**Authors:** Yingying Wang, Scott K. Holland

## Abstract

Magnetoencephalography (MEG) records brain activity with excellent temporal and good spatial resolution, while functional magnetic resonance imaging (fMRI) offers good temporal and excellent spatial resolution. The aim of this study is to implement a Bayesian framework to use fMRI data as spatial priors for MEG inverse solutions. We used simulated MEG data with both evoked and induced activity and experimental MEG data from sixteen participants to examine the effectiveness of using fMRI spatial priors in MEG source reconstruction. Our results provide empirical evidence that the use of fMRI spatial priors improves the accuracy of MEG source reconstruction.

## Introduction

The last decade has witnessed great advances in multi-modal data fusion techniques (Auranen et al., 2009; Baillet & Garnero, 1997; Baillet, Garnero, Marin, & Hugonin, 1999; Debener, Ullsperger, Siegel, & Engel, 2006; Friston et al., 2008; Henson, Flandin, Friston, & Mattout, 2010; Nummenmaa et al., 2007; Sato et al., 2004; Wipf & Nagarajan, 2009) and increasing interests in studying high-order cognition in the human brain using multi-modal techniques, especially data fusion of functional magnetic resonance imaging (fMRI) and magnetoencephalography (MEG) (Liljestrom, Hulten, Parkkonen, & Salmelin, 2009; Pang, Wang, Malone, Kadis, & Donner, 2010; Vartiainen, Liljestrom, Koskinen, Renvall, & Salmelin, 2011; Wang, Holland, & Vannest, 2012). Our previous study has demonstrated spatial concordance in the left inferior frontal gyrus (IFG) for covert or overt generation versus overt repetition, and bilateral motor cortex when overt generation versus covert generation (Wang et al., 2012), algin with other studies (Pang et al., 2010; Vartiainen et al., 2011) and provide evidence that the two modalities are assessing the same language network during language production and comprehension tasks. The high spatial concordance between fMRI and MEG data in accessing language function provides confidence that fMRI can be used as spatial constraints on MEG source localization. Consequently, high spatiotemporal information obtained from the promising multi-modal data integration approach can potentially yield new insights into the complex brain networks supporting high-order cognition. Several integration schemes have been introduced to combine fMRI and MEG data such as fMRI-guided equivalent current dipole (ECD) fitting (Ahlfors & Simpson, 2004), fMRI-constrained cortical current density imaging (Dale et al., 2000; Liu et al., 2008; Ou et al., 2010), and the recent popularity of Bayesian schemes applied in MEG source inversion (Auranen et al., 2009; Friston et al., 2008; Henson et al., 2010; Wipf & Nagarajan, 2009).

Currently there is no best solution for combining fMRI and MEG data, but only most suitable approach to integrate the two modalities based on the neurobiological characteristics of brain functions. Based on the current literature, parametric empirical Bayesian (PEB) (Henson et al., 2010) with distributed cortical current density imaging approach is well suited to high-order cognitive data on several counts. First, language tasks usually require communication and coordination among many regions distributed in the brain. It is hard to account for highly distributed regions involved in language tasks using ECD modeling which represents a relatively small number of focal sources. Second, researchers (Liljestrom et al., 2009; Vartiainen et al., 2011; Wang et al., 2012) have shown that there is no simple one-to-one spatial correspondence between fMRI and MEG results. For example, high concordance between MEG and fMRI was observed in the left inferior frontal gyrus (IFG), the bilateral motor cortex, and the right insula, but not in the left medial frontal gyrus and the left cingulate gyrus during an overt verb generation task (Wang et al., 2012). The PEB approach allows the use of fMRI spatial information as “soft” constraints rather than “hard” ones. Alas, current available integration schemes have only been tested in the experimental paradigms with short-onset and time-locked stimuli such as visual, auditory, motor, somatosensory. Note it is easy to design a short paradigm with hundreds of trials consisting of these stimuli so that high SNR can be achieved for ECD fitting by averaging hundreds of trials of external MEG signals. Thus, for these types of paradigm design, incorporating fMRI spatial information into MEG source inversion offers very limited advantage and is not a cost-effective approach since MEG alone can already achieve sufficient localization accuracy with hundreds or even thousands of trials of neuromagnetic data. However, for high-order cognitive tasks, especially natural language processes like story listening (narrative comprehension), it is extremely hard to acquire hundreds of trials within a short time period. Further, it is difficult to separate evoked and induced responses to the stimulus, complicating the source localization process from MEG data alone. Long time intervals can potentially induce fatigue and more head motion which deteriorate the quality of neuroimaging data. Therefore, for these types of cognitive tasks, the extra information from fMRI can play a crucial role in improving the solutions of ill-posed MEG source inversion and add another perspective to our understanding of the neurobiology of language.

In this study, we elaborate on the theory of PEB approach and provide information on integration of fMRI spatial constrains into MEG source localization. We present empirical evidence using simulations and analysis of experimental data from sixteen participants who performed a narrative comprehension task during MEG recordings.

## Methods

### The MEG Inverse Problem

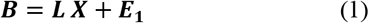

***B*** is the vector of magnetic signal at a given time sample, ***L*** is the lead-field matrix, ***X*** is the unknow brain activities, ***E*_1_** is assumed to be Gaussian noise with zero mean and known variance. The linear system presented in (1) is underdetermined since the number of sensors (only hundreds) is much fewer than the number of possible sources (over thousands). In order to find solutions for (1), there are two general methods including widely used ordinary least squares estimation (OLSE) and maximum likelihood estimation (MLE). The former requires no assumptions about the distributions, but does not allow model testing or selection, etc. On the contrary, MLE provides complete and consistent solutions and can serve as a steppingstone for other inference methods such as Bayesian methods, inference with missing data, etc. We will formulate the inversion of (1) within a probabilistic Bayesian framework which allows us to choose hyperparameters and optimize MEG source inversion by incorporating fMRI spatial information.

### Bayesian Framework

The probabilistic model for sources ***X*** can be expressed under the assumption of Gaussian noise at the sensor level. Then, the probability distribution is given by (2).

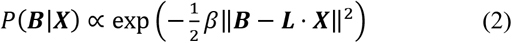

*β* denotes the noise variance. The magnetic field ***B*** is observed for a given source current ***X*** is proportional to Gaussian probability density. According to Bayes’ theorem, (2) can be used to form the posterior probability over unknown ***X*** by (3).

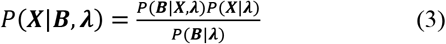

***λ*** represents all the other assumptions and beliefs about the model, the denominator in (3) is the evidence for ***λ*** and ensures the posterior probability is normalized.

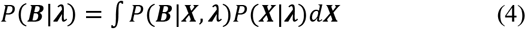

The prior probability density *P* (***X|λ***) represents all the he prior information known about the unknown ***X*** and constrains before data is seen and can be regarded as a regularization which limits model overfitting. Since *P*(***B|λ***) does not depend on ***X***. It can be omitted to generate the unnormalized posterior density in (5).

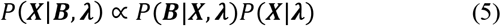

The posterior probability is proportional to likelihood times the prior probability. Possible solution of ***X*** must simultaneously give a high data likelihood *P*(***B|X, λ***) and be probable under constraint of the *prior P*(***X|λ***) to give an appreciable posterior distribution *P*(***X|B, λ***). The posterior probability shown in (3) is a measure of what is known after the data is seen and quantifies any new knowledge gained. The data likelihood *P*(***B|X, λ***) is a measure of how well the model predicted the data and essentially determines whether the data under investigation contains any new information. Bayes’ theorem enables us to update the distribution over parameters from the prior to the posterior distribution over the latent variables in light of observed data. For a simple illustration, assume that non-informative priors are used. For each voxel in the brain, we believe there is a 50 percent chance it will be active or inactive. Then, via Bayesian statistics, we update our old beliefs iteratively so some voxels might have 90 percent chance to be active during the task based on the data and priors.

Using the expectation maximization (EM) algorithm, the model is learned by alternating between estimating the posterior distribution over latent variables for a particular setting of model parameters and then re-estimating the best-fit parameters given that distribution over the latent variables (Beal, 2003; Friston, Mattout, Trujillo-Barreto, Ashburner, & Penny, 2007). In Bayesian statistics, maximum a posteriori probability (MAP) can be used to as a point estimate to approximate the posterior distribution. There are four major ways of computing MAP estimates, depending upon the specifics of the problem. First, when conjugate priors are used, MAP estimates can be solved analytically since the mode of the posterior distribution can be given in closed form. Second, MAP estimates can be computed analytically or numerically through numerical optimization such as conjugate gradient method or Newton’s method, which requires first or second derivatives. Third, MAP estimates can be obtained via a modification of an EM algorithm which does not require derivatives of the posterior density. At last, MAP can also be calculated through a Monte Carlo method using simulated annealing.

### PEB Framework

In PEB framework (Henson et al., 2010), the third type of method is used to obtain MAP estimates via variational free energy under the Laplace approximation (Friston et al., 2007). Variational Bayes (VB) by variational free energy is a generic approach to construct an analytical approximation of the posterior probability distribution (Bishop, 1998; Friston et al., 2007). The foundation of VB is that the log-evidence can be defined in terms of the free energy *F* and a Kullback-Leibler divergence term.

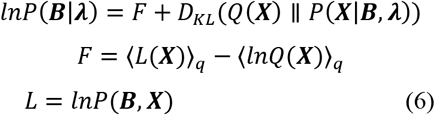

〈*L*(*X*)〉_*q*_ is the expected energy. 〈*lnQ*(***X***)〉_*q*_ is the entropy measuring the uncertainty in a random variable in information theory. From (2), the free energy is a lower-bound approximation to the log-evidence because the divergence term is always positive. The goal is to compute *Q*(***X***) for each model by maximizing the free energy *F*. Then, compute *F* for Bayesian inference and model comparison, respectively.

### FMRI Priors

FMRI results are usually presented as noise-normalized statistical parametric maps (SPMs) rather than as maps of raw signal strength since noise variance may vary greatly between voxels. The topological features of these fMRI SPMs can be assigned probabilities that quantify the chance that the cluster of voxels can be active under null hypothesis. A voxel with the increases in the mean fMRI signal associated with one experimental condition versus another indicates that it has high probability to be an “active” source in MEG source space. Thus, fMRI results can be projected onto the cortical mesh and then converted into covariance matrices (Henson et al., 2010). Since the underlying physiological connections between fMRI and MEG signals are still unclear, fMRI data are treated as probabilistic information about the spatial location of active regions in the brain rather than quantitative information about the amplitude of the neural activity. First, we define a number of discrete clusters by thresholding the SPMs since using each suprathreshold “cluster” from fMRI SPMs to form a separate prior rather than a single prior enables flexibility in adjusting different contributions of multiple constrains. Second, the fMRI clusters need to be projected onto the cortical mesh since the fMRI clusters are 3D volume while MEG cortical mesh is based on the cortical surface through (7). 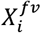 is the *i* th fMRI cluster. 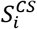 is its counterpart on the cortical surface. *ε_i_* is the error term and *f* is the linkage function that is Heaviside function binarizing each fMRI spatial prior. *H* is the Voronoi-based interpolation function (Kiebel, Goebel, & Friston, 2000) that has been suggested to be more superior than other interpolation methods (Henson et al., 2010).

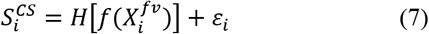

At last, these cortical patches need to be transferred to covariance components through (8). Before conversion, an extra spatial smoothing step is added to reduce the misregistration errors via a spatial coherency function.

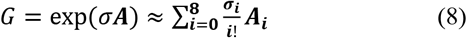

The elements *a_ij_* in matrix **A** equals 1 if *i* and *j* are neighbors and 0 otherwise. *σ* is the smoothing parameter. After smoothing 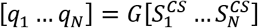, the covariance components can be generated by the outer product 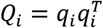.

### Data Simulation

In (1), **L** was computed using a single sphere head model (Sarvas, 1987). Two dipoles at *X*_1_(−38, 43, 5) and *X*_2_(−54, – 13, 5) in Montreal Neurological Institute (MNI) space were used to generate the simulated MEG data **B**. In order to check how *priors* affect both evoked and induced brain signals, *X*_1_ and *X*_2_ had different time courses in (9).

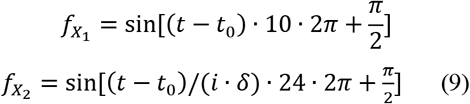

In (9), *t* ∈ [0.55,0.85] denotes time in seconds, *i* ∈ [1,100] denotes the trial number, *δ* is a positive random number that is different for each trial so that *X*_2_(−54, −13, 5) contains induced activity. We also added gaussian noise to the simulated MEG data to generate data with three different noise levels including no noise, signal-to-noise ratio = 10 *dB* and −10 *dB.* The MATLAB code for simulating MEG data is available by request and the illustration diagram of MEG data simulation process is in supplementary materials.

The fMRI SPM{T} map was generated using MarsBaR (Brett, Anton, Valabregue, & Poline, 2002) and then projected to the cortical mesh using SPM8 (www.fil.ion.ucl.ac.uk/spm/). Both valid and invalid fMRI spatial *priors* were considered so that we can evaluate the impact of invalid *priors* that mismatch MEG dipole locations (see Fig. 1).

**Fig. 1.**
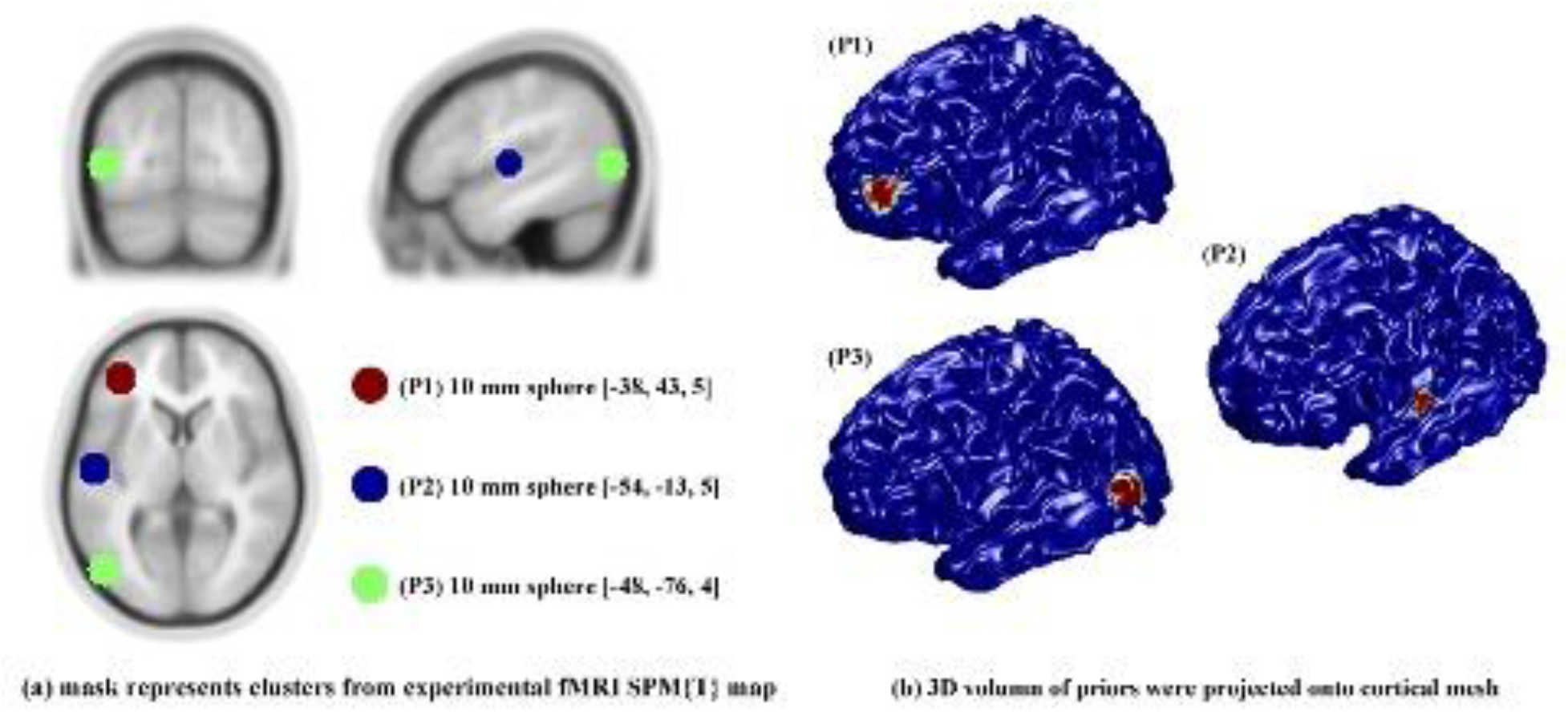
Hypothetic fMRI priors: (a) mask represents clusters from experimental fMRI SPM{T} map superimposed on a template MRI. (b) 3D volumes of priors were projected onto the cortical mesh.

For evaluation of source reconstruction results, we used absolute measure of localization error denoted by Euclidean distance between the estimated peak location and the actual location of the sources. We also used area under curve (AUC) based on receiver operating characteristic (ROC) curve to quantify the detection accuracy of various inverse solutions (Barnes, Chowdhury, Lina, Kobayashi, & Grova, 2013; Grova et al., 2006). The simulated MEG data provides the ground truth. The various inverse solutions (no fMRI *priors*, with all fMRI *priors*, with only valid fMRI *priors,* with only invalid fMRI *prior)* need a decision threshold *β* to build ROC curves. By comparing the inverse solutions with the gold standard, we were able to quantify the number voxels detected as true positive (TP), true negative (TN), false positive (FP) and false negative (FN) for each threshold *β* ∈ [0,1]. Sensitivity and specificity were then estimated using (10).

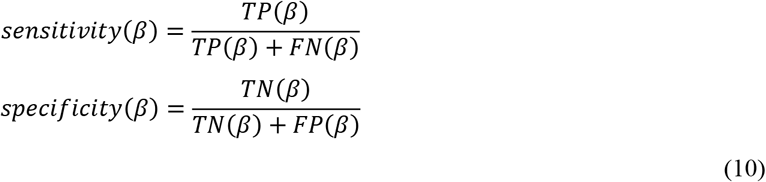

ROC curves were plotted using *sensitivity(β)* as y-axis and *(1-specificity(β))* as x-axis for each threshold *β* ∈ [0,1]. Then, AUC was computed to evaluate detection accuracy. In general, an AUC value greater than 0.8 is considered as sufficiently accurate that achieves 80% good detection rate. Higher AUC indicates better performance. For evaluation of time-course results, we used scatter plots to visualize the correlation between the “true” time courses and the estimated time courses in both time and frequency domain. The volume of interests had a radius of 10 mm. Time courses were extracted from the first eigenvariate of all voxels in the volume of interests, rather than the mean values, since the eigenvariate value is more robust to heterogeneity of response within a cluster. The root mean square error (RMSE) were computed between the “true” time courses and the estimated time courses in both time and frequency domain.

### Experimental MEG Data

The experimental MEG and fMRI data from sixteen participants with average age 15.8 years were described in our previous study and a detailed description of the data and paradigms can be found in (Wang et al., 2012). MEG data were acquired using a 275-channel whole head MEG system (VSM Med-Tech Ltd., Port Coquitlam, BC, Canada) sampled at 6 KHz and fMRI data were acquired on a Philips Achieva 3-Tesla MRI scanner with Dual Quasar gradients (Philips Medical Systems, Best, The Netherlands). During the scan, the participants performed a narrative comprehension task including three conditions (story listening, question answering, and pure tone listening). ^1^ In the current study, we only focused on the contrast of story listening versus tone listening. The group fMRI results were used as spatial priors for MEG source reconstructions (see Fig. 2). Three clusters that survived the thresholding (height threshold *T* = 5.78 and extent threshold *k* = 50 voxels, *p* < 0.005 Family-Wise Error rate corrected) were projected onto the template cortical mesh in Fig. 2–1 and correspond approximately to left inferior gyrus (IFG) and bilateral superior temporal gyrus (STG).

**Fig. 2.**
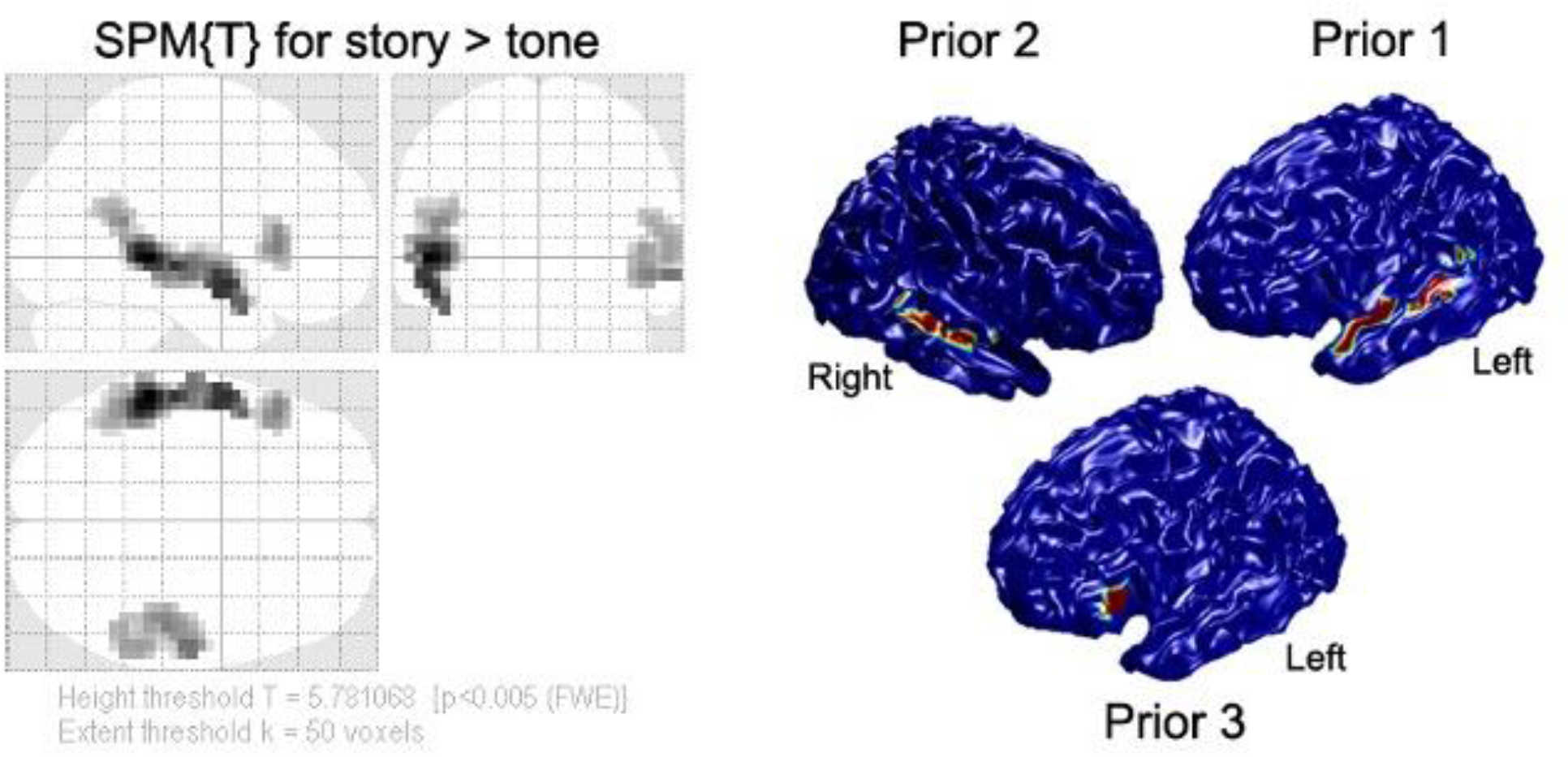
Threshold SPM{T} for group fMRI results in MNI standard space from 16 participants. There priors are generated from group fMRI results and projected on the MNI template cortical mesh. Color reflects T-value (scale irrelevant other than blue regions having value zero).

## Results

### Simulation Study

Table 1 summarized the location error for both peak and center of mass from different inversion methods including multiple sparse priors (MSP) without spatial priors, MSP with all priors, MSP with only valid priors, and MSP with only invalid priors (see supplementary Fig. S2), as well as the AUC values. The ROC curves were plotted in Fig. 3.

**Table 1.** Summary of different inversion methods

| Localization Error (mm) |  | No priors | All | Valid Only | Invalid Only |
| --- | --- | --- | --- | --- | --- |
| No noise |  |  |  |  |  |
| $X_1$ | Peak | 9.70 | 9.70 | 9.70 | 9.70 |
|  | Cmass | 8.15 | 7.80 | 7.80 | 8.15 |
| $X_2$ | Peak | 7.35 | 5.83 | 5.83 | 7.35 |
|  | Cmass | 6.75 | 1.56 | 1.56 | 6.75 |
| AUC |  | 0.82 | 0.99 | 0.99 | 0.85 |
| Signal-to-Noise Ratio = 10 dB |  |  |  |  |  |
| $X_1$ | Peak | 9.70 | 9.70 | 9.70 | 9.70 |
|  | Cmass | 8.15 | 7.80 | 7.80 | 8.15 |
| $X_2$ | Peak | 7.35 | 5.83 | 5.83 | 7.35 |
|  | Cmass | 7.10 | 1.64 | 1.64 | 7.10 |
| AUC |  | 0.82 | 0.99 | 0.99 | 0.85 |
| Signal-to-Noise Ratio = -10 dB |  |  |  |  |  |
| $X_1$ | Peak | 9.70 | 9.70 | 9.70 | 9.70 |
|  | Cmass | 8.76 | 8.50 | 8.50 | 8.76 |
| $X_2$ | Peak | 7.35 | 5.83 | 5.83 | 7.35 |
|  | Cmass | 7.49 | 1.81 | 1.81 | 7.53 |
| AUC |  | 0.80 | 0.95 | 0.96 | 0.84 |
| Cmass: the center of the cluster |  |  |  |  |  |

**Fig. 3.**
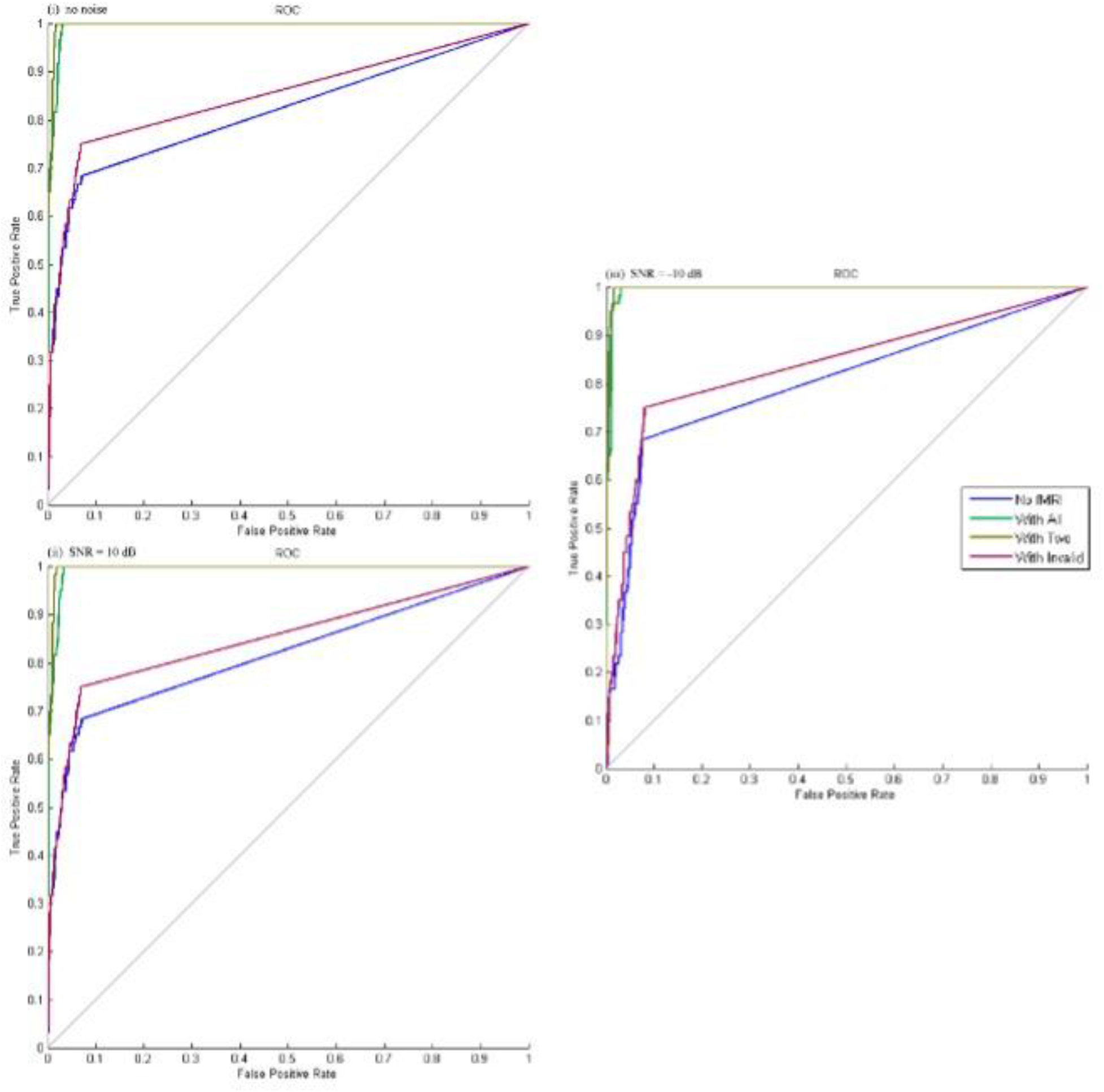
The ROC curves for different inversion methods. Blue: Inversion without fMRI priors; Green: Inversion with all fMRI priors including valid and invalid; Khaki: Inversion only with valid fMRI priors; Red: Inversion only with invalid priors.

Note that the ROC curves had a non-smooth appearance due to the small number of true positive voxels embedded in the simulated data set relative to true negative voxels. True positive voxels were detected as positive at an abrupt threshold, the distribution of values was not very wide ranging. Still the AUC varies with the prior information incorporated in the model and from this parameter we can see that the performance is superior when valid fMRI priors or mixture of valid and invalid fMRI priors used in the estimation process.

For evoked source *X*_1_(−38, 43, 5), the location error of peak activity was not affected by the fMRI priors, but the location error of the center of cluster decreased when the accurate fMRI priors were incorporated. In addition, the mixture of spatial priors with valid and invalid locations decreased the location error of the center of cluster. The AUC increases from good to excellent when the accurate fMRI priors were used.

For induced source *X*_2_(−54, −13, 5), the location errors of peak activity and the center of cluster were greatly reduced by the fMRI priors. Higher-order cognitive activity in the brain has been shown to be induced responses. Even when only inaccurate spatial prior was used, the location error and AUC was not affected that much. The add-on white noise slightly decreased the AUC and increased the location error for both sources.

The extracted time courses were plotted against the actual time courses (see supplementary Fig. S6–7). Table 2 showed the RMSE values for different inversion methods.

**Table 2.** RMSE as a quantitative measure of time-course extraction quality for each inversion method.

| RMSE | No priors | All | Valid Only | Invalid Only | Virtual Sensor |
| --- | --- | --- | --- | --- | --- |
| $X_1$ | | | | | |
| Average of 100 trials | 0.015 | 0.001 | 0.004 | 0.011 | 0.007 |
| 100 trials | 0.020 | 0.002 | 0.005 | 0.016 | 0.010 |
| $X_2$ | | | | | |
| Average of 100 trials | 0.033 | 0.024 | 0.037 | 0.021 | 0.008 |
| 100 trials | 0.490 | 0.494 | 0.488 | 0.497 | 0.505 |

The introduction of spatial priors reduced discrepancy between the reconstructed time courses and ground truth, which were more evident in evoked source activity *X1*. For evoked source activity *X*_1_, average of 100 trials did not change the RMSE compared to 100-trial time courses. But for induced source activity *X*_2_, average of 100 trials reduced the RMSE dramatically compared to 100-trial time courses. Inversion with all fMRI spatial priors including invalid and valid ones offered the lowest RMSE for evoked source activity *X*_1_, while inversion with valid only spatial fMRI priors gives lowest RMSE for induced source activity *X*_2_ without averaging. The virtual sensor approach gave the lowest RMSE for induced source activity *X*_2_ after averaging all the trials. The inversion approach with all fMRI spatial priors provides the lowest overall RMSE for both evoked and induced source activity (*X*_1_ and *X*_2_). The time course extraction was problematic for induced response *X*_2_ due to the trial-to-trial variability. The unaveraged time courses had higher RMSE which indicated high discrepancy between the original signal and extracted signal.

The single-sided amplitude spectrum plots were generated for all time course using Fourier transformation (see supplementary Fig. S8). All inversion methods yielded low RMSE. Table 3 showed the RMSE for spectrum plots.

**Table 3.** RMSE as a quantitative measure of time-course extraction quality for each inversion method.

| RMSE | No priors | All | Valid Only | Invalid Only | Virtual Sensor |
| --- | --- | --- | --- | --- | --- |
| $X_1$ | | | | | |
| Average of 100 trials | 0.0009 | 0.0001 | 0.0002 | 0.0007 | 0.0004 |
| 100 trials | 0.0010 | 0.0001 | 0.0003 | 0.0008 | 0.0005 |
| $X_2$ | | | | | |
| Average of 100 trials | 0.0016 | 0.0011 | 0.0019 | 0.0009 | 0.0004 |
| 100 trials | 0.0026 | 0.0022 | 0.0029 | 0.0021 | 0.0017 |

### Experimental MEG Data

The introduction of group fMRI results as spatial priors for MEG inverse problem significantly increased free-energy *F* (*T*(15) = 2.51, *p* < 0.05, one-tailed). For evoked activity, the effect of fMRI priors on the source reconstruction was mainly to divide activity slightly more bilaterally in the STG and reduce the spurious clusters (see Fig. 4). For induced activity, the effect of fMRI priors on the source reconstruction was to pull activity slightly more left in STG (see Fig. 4) that is consistent with known patterns of leftward lateralization of this task (Wang et al., 2012).

**Fig. 4.**
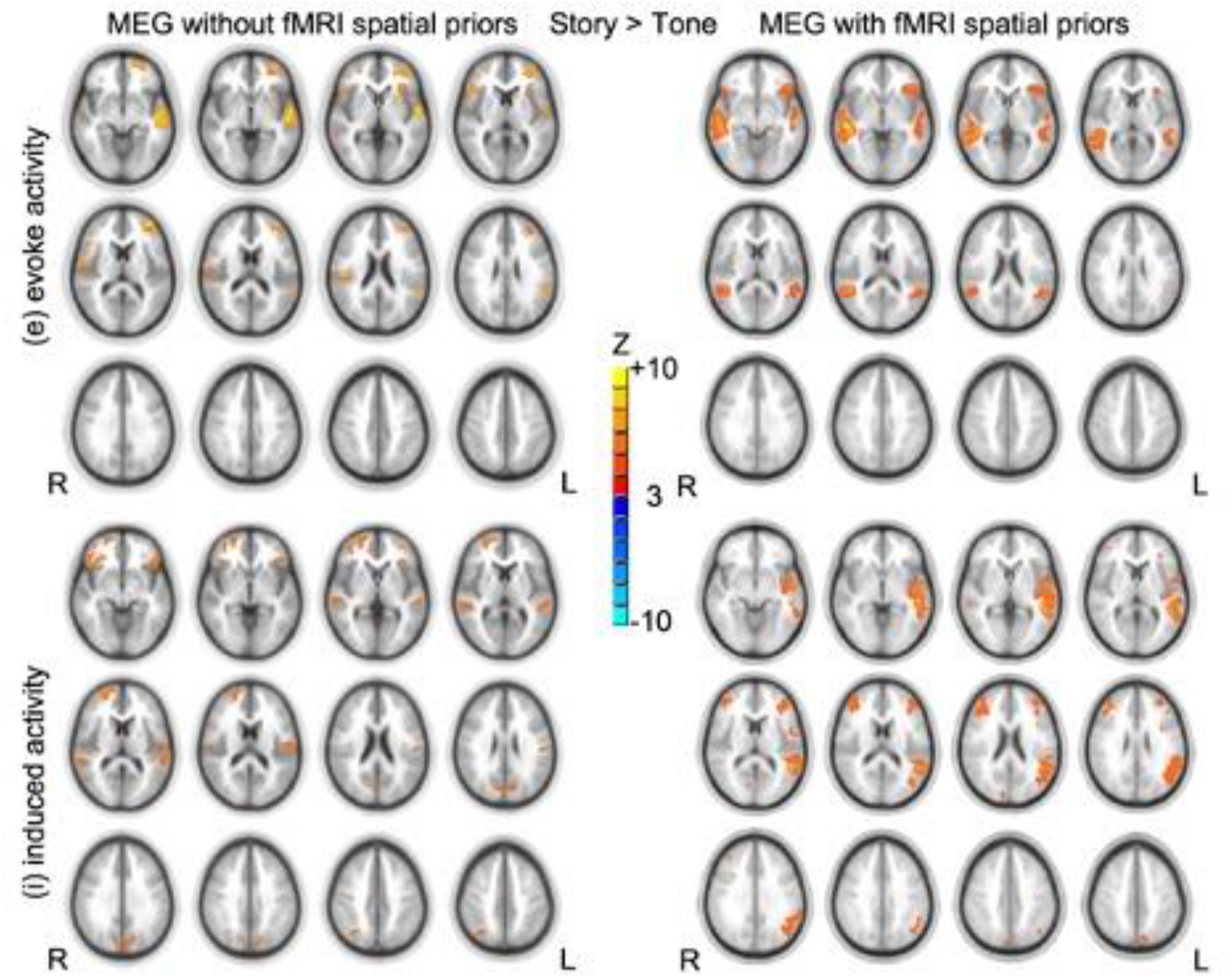
Group composite activation maps of story > tone contrast from MEG without fMRI spatial priors (left) and MEG with fMRI spatial priors (right) results (N = 16). Significant level: p < 0.05 false discovery rate corrected. Cluster size > 20. Slice range: Z = −5 to +50 mm (Talairach coordinates) and 5 mm between each successive slice displayed. All images are in radiologic orientation (left on the right, right on the left).

## Discussion

We implemented the hierarchical Bayesian framework on MEG source reconstruction with fMRI spatial information incorporated as spatial priors and applied this approach to both simulated MEG data and experimental MEG data from sixteen adolescents during a narrative comprehension task.

For simulated MEG data, the spatial resolution of MEG source reconstruction increases (3 mm on average) by incorporating the prior information from fMRI in the source reconstruction. The use of fMRI spatial priors greatly reduced location error for induced source in MEG data. This is important since the induced responses of the neuronal activity in the brain are often corresponding to high-order cognitive tasks. Therefore, the additional spatial priors from fMRI could benefit the accuracy of source reconstruction especially for this type of high-order cognitive processes. Wang et al. (2012) reported similarities and differences between fMRI and MEG data from the same participant (Wang et al., 2012). FMRI spatial priors would include valid and invalid spatial locations. Our results suggested that the combination of accurate and inaccurate spatial priors still increased the accuracy of MEG source reconstruction since the inaccurate priors are effectively discarded in the restricted maximum likelihood procedure. The AUC was greatly increased from good to excellent when the fMRI priors are incorporated into the MEG inversion problem. Thus, the MSP with fMRI priors is a robust approach for MEG inverse problem.

This is the first study that applied the hierarchical Bayesian framework on simulated MEG data from both evoked and induced source activity. The evoked responses are phase-locked to trial onset and induced responses have a random phase-relationship over trials. Simple paradigm design with short duration of visual, or auditory, or sensory, or motor stimulus usually produces evoked responses. High-order cognitive paradigms have longer stimulus duration and induce a more complex process in the brain that varies over trials. This type of paradigm usually generates both evoked and induced activity in the brain. From our simulated data, we found that the accurate fMRI spatial priors have evident effects on the induced activity not the evoked activity. The induced responses benefit more from the fMRI prior information since there is more trial-to-trial variability in the induced responses. From our experimental data, activation patterns from evoked and induced responses separate different stages of language processes involved in the narrative comprehension task. This finding demonstrates that the wide range of neuronal activity recorded by MEG could improve our understanding of language processes.

The introduction of accurate spatial priors reduced discrepancy between the reconstructed time courses and the actual time courses. The effect was more evident in the estimated time courses of the evoked source. This finding is encouraging in that spatial priors are beneficial not only in spatial accuracy but also in temporal accuracy. The trial-to-trial variability for induced source affected the accuracy of estimated time courses at each individual trial due to random nature of the phase relationship with the stimuli. When the time courses from induced source were transferred into frequency domain, the power spectrum in the frequency domain for each trial showed much less discrepancy between the ground truth and estimation. This finding confirmed other studies in the literature (David, Kilner, & Friston, 2006; Friston, 2006; Michalopoulos, Iordanidou, Giannakakis, Nikita, & Zervakis, 2011), suggesting using the average energy over trials to represent the induced responses.

For our experimental data, the introduction of fMRI spatial priors showed significant increases in the free-energy bound, which indicates the improvement in model evidence by adding fMRI spatial information into MEG source reconstruction. Our experimental results are in line with Henson et al. (Henson, Wakeman, Litvak, & Friston, 2011). The group composite activation maps of story > tone contrast from MEG with fMRI spatial priors showed very similar activation patterns to our previous cross-sectional fMRI studies (Karunanayaka et al., 2007; Szaflarski et al., 2012; Vannest et al., 2009). FMRI spatial information reduced the spurious clusters for evoked activity and showed more left-lateralized activation pattern for induced activity. The activations in the STG were bilateral for evoked activity, whereas activations in the STG were left-lateralized for induced activity. The bilateral activations in STG, revealed by evoked responses, could be due to the residual prelinguistic auditory processing since the use of tone listening as the control condition presumably subtracts out earlier stages of auditory processing. The strong left-lateralized activations in STG, revealed by induced responses, agree with other fMRI studies (Benson, Richardson, Whalen, & Lai, 2006; Binder et al., 2000; Mummery, Ashburner, Scott, & Wise, 1999; Scott, Blank, Rosen, & Wise, 2000), which suggested the high-order cognitive process for understanding speech sounds takes place in the left temporal lobe. Therefore, MEG with fMRI spatial priors might be helpful in determining the lateralization of language functions in presurgical mapping.

## Conclusion

In summary, the hierarchical Bayesian framework allows us to incorporate fMRI spatial priors and improve MEG source estimates resulting in a distribution of likely solutions instead of a single solution. Combining MEG and fMRI data within the Bayesian framework is a promising approach to overcome the limitations of each modality and offers brain signal with fine spatiotemporal resolution which is crucial for effective connectivity analysis.

## Acknowledgments

The authors would like to thank Ms. Kate Hibbard, Ms. Julie Franks, Ms. Sara Robertson, and Ms. Amanda Huber for their assistance in helping with recruitment and data collection, as well as Mr. Kendall O’Brien and Ms. Amanda Woods for their assistance in performing all the MRI scans.

## Declarations

### Funding

This work was supported by the U.S. National Institute of Child Health and Human Development under Grant R01-HD38578 awarded to S.K. Holland.

### Conflicts of interest

Both authors declare no competing interests.

### Ethics approval

All procedures performed in studies involving human participants were in accordance with the ethical standards of the Institutional Review Board of Cincinnati Children’s Hospital Medical Center.

### Informed consent

Informed consent was obtained from all participants.

### Consent for publication

Both authors consent for publication.

### Availability of data and material

All data are available by request.

### Code availability

All codes are available by request.

### Authors’ contributions

Author contributions included conception and study design (YW and SKH), data collection and analysis (YW), interpretation of results (YW and SKH), drafting the manuscript work or revising it critically for important intellectual content (YW and SKH) and approval of final version to be published and agreement to be accountable for the integrity and accuracy of all aspects of the work (Both authors).

## Supplementary materials

**Fig. S1.**
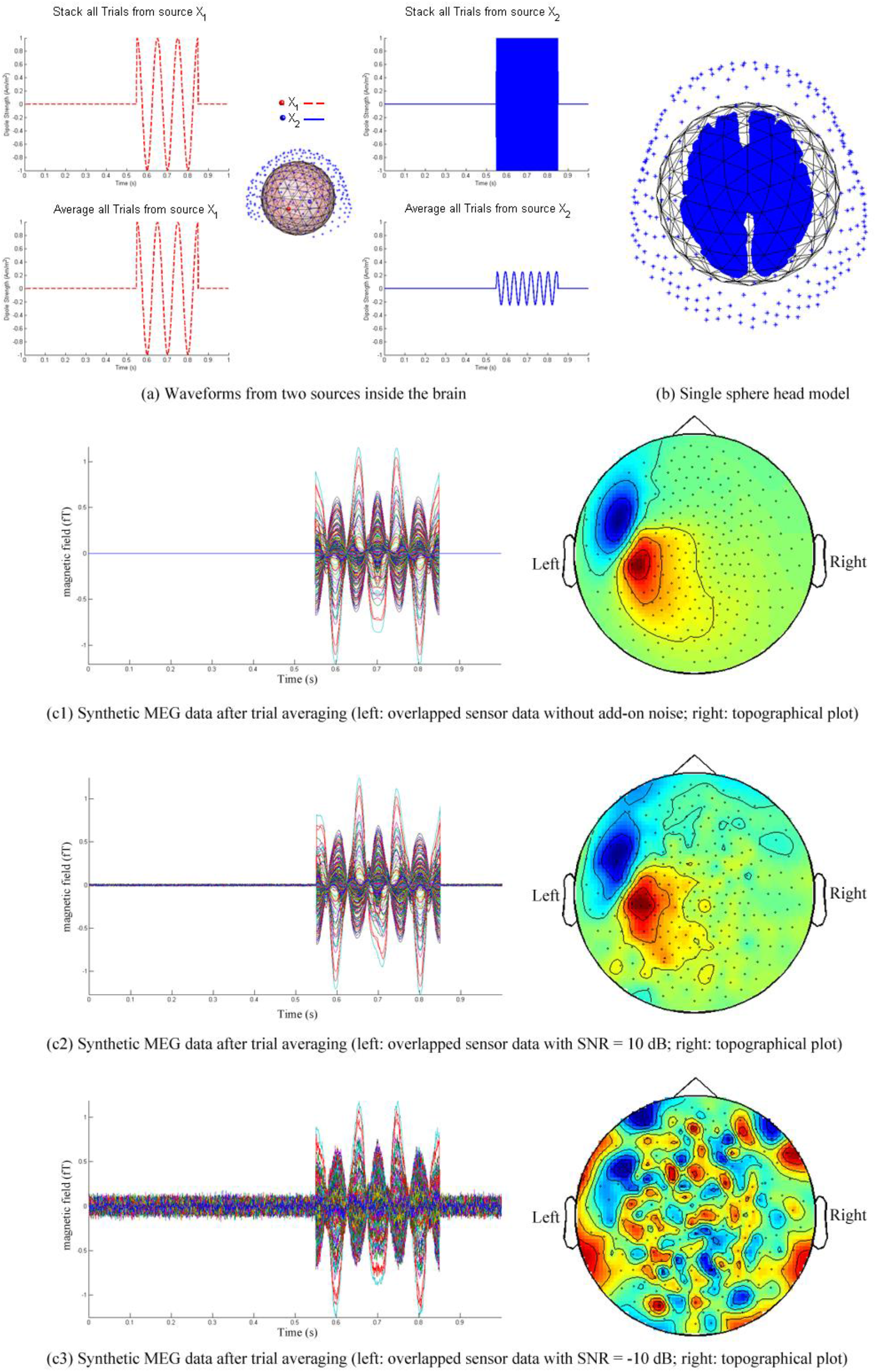
Illustration diagram showing MEG data simulation process

**Fig. S2.**
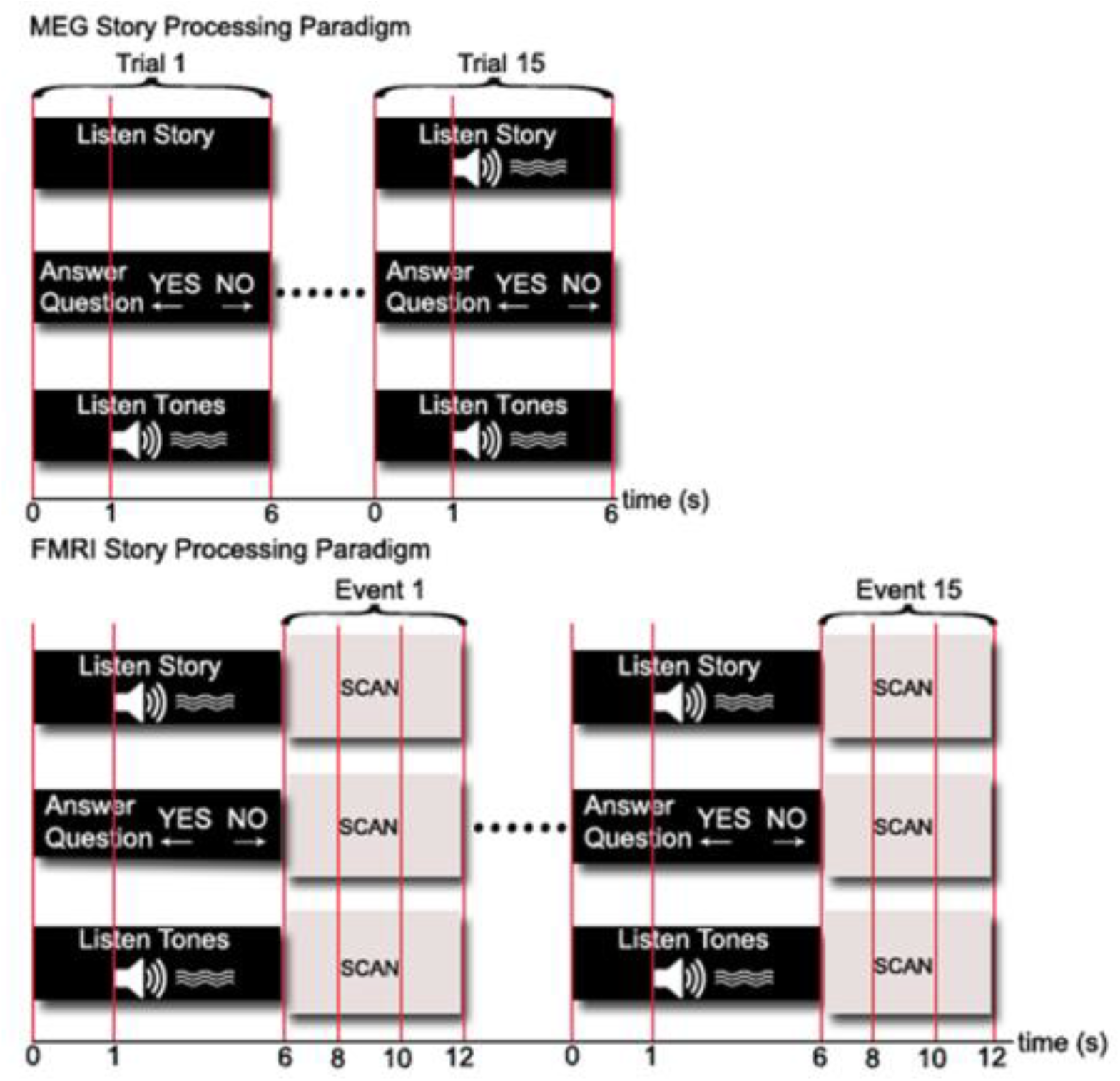
Timing diagram of the event-related story-processing task for MEG and fMRI task. MEG: 15 5-s Trials for each condition are recorded. FMRI: 15 36-s cycles for each phase of the paradigm are presented for a total scan time of 9 minutes. Online performance data of correctness and response time are recorded for both MEG and fMRI sessions.

**Fig. S3.**
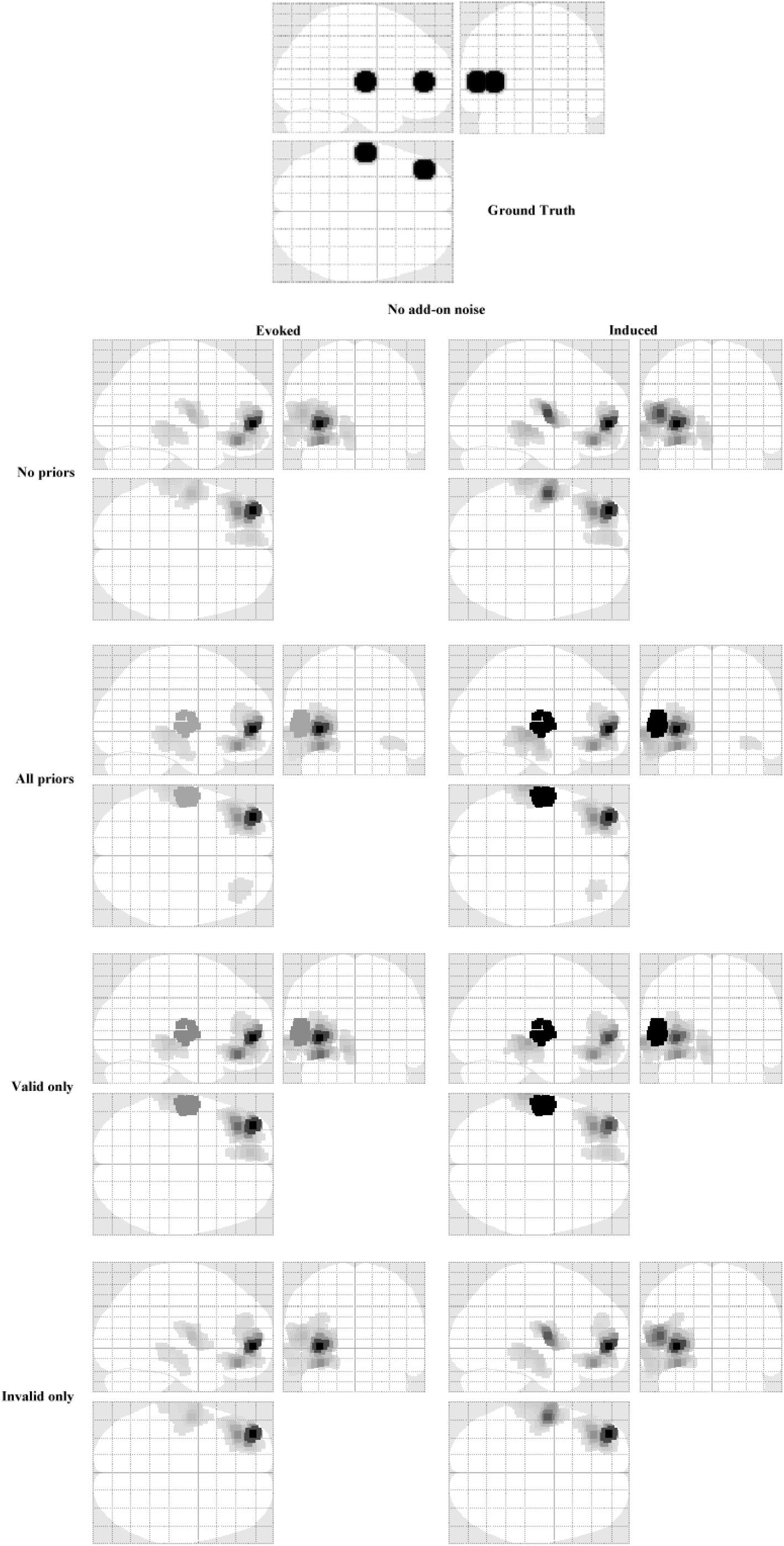
No add-on noise: source reconstruction results for four different inversion schemes including no priors, all priors, valid only, and invalid only.

**Fig. S4.**
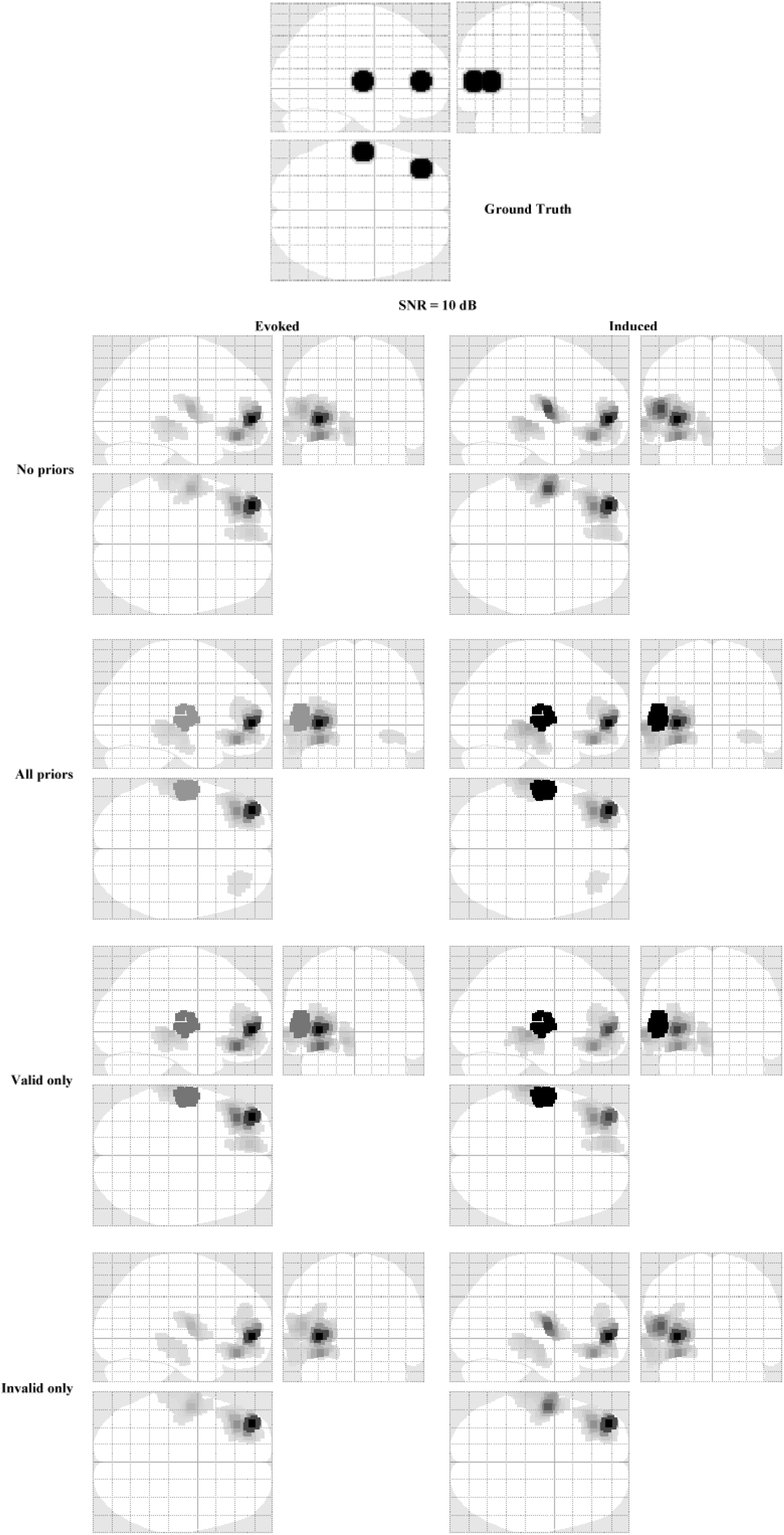
SNR = 10 dB: source reconstruction results for four different inversion schemes including no priors, all priors, valid only, and invalid only.

**Fig. S5.**
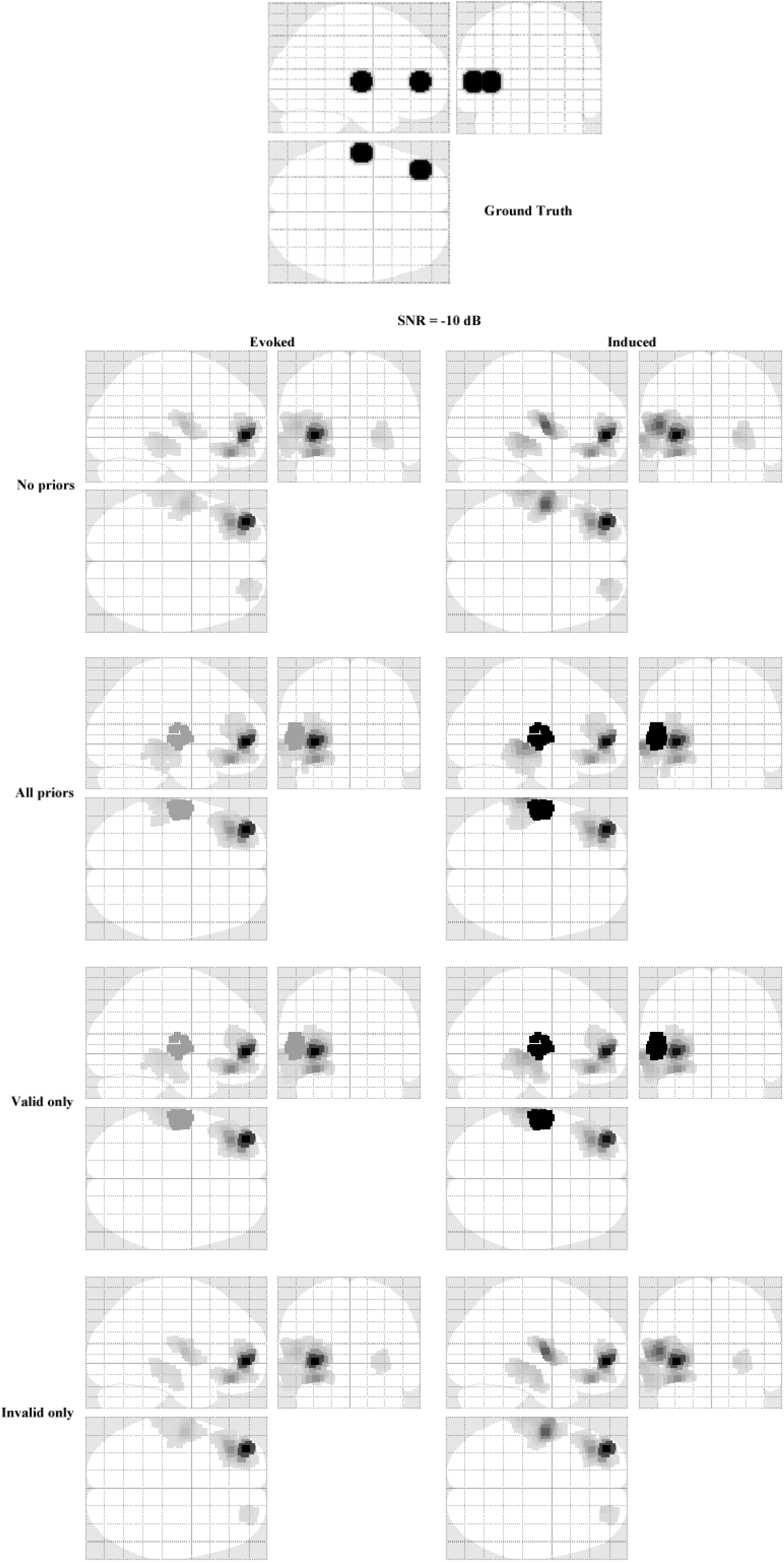
SNR = −10 dB: source reconstruction results for four different inversion schemes including no priors, all priors, valid only, and invalid only

**Fig. S6.**
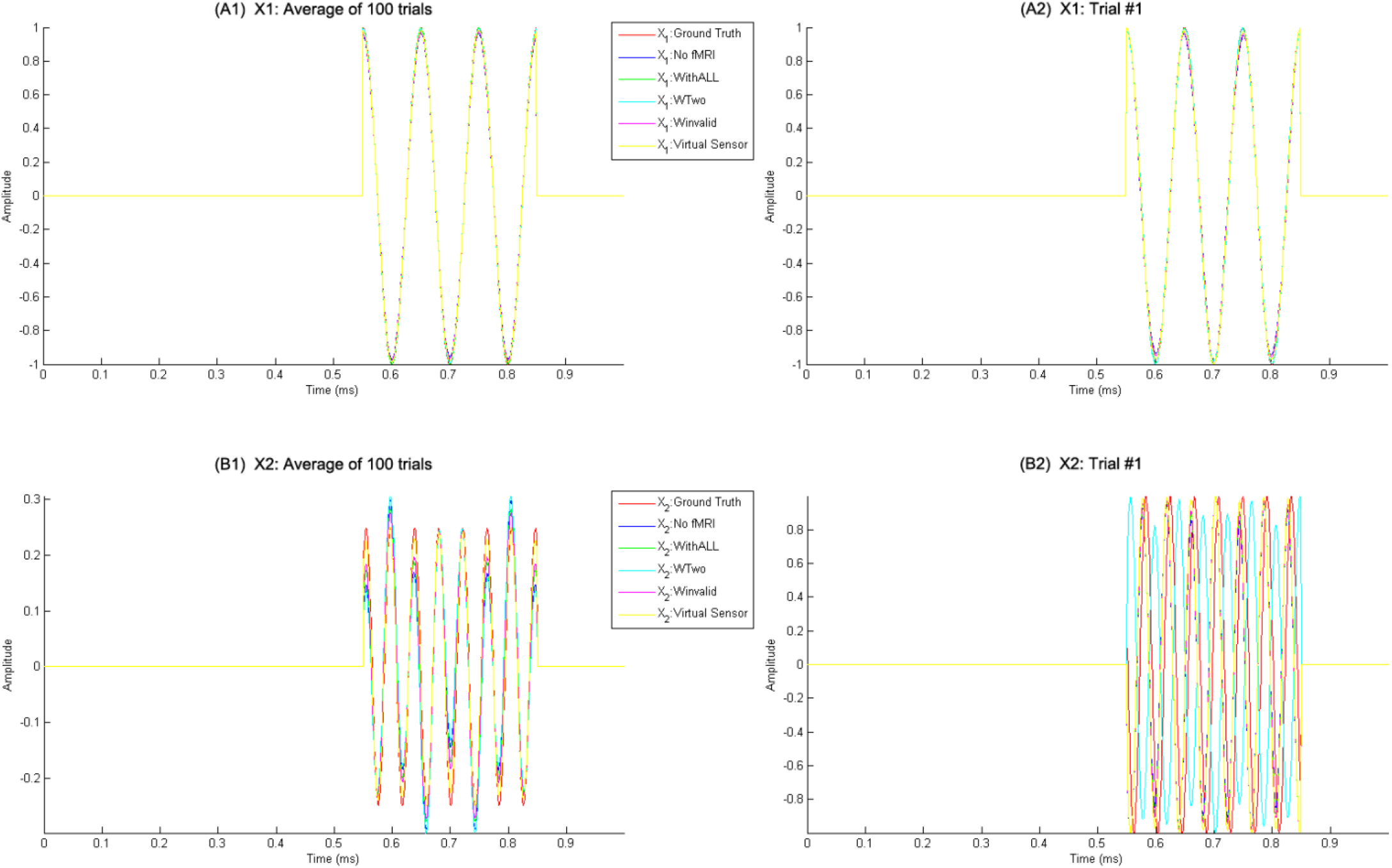
Time course plots: (A1): 100 trials for source *X1* (−36, 52, 2) were averaged and color coded by the type of inversion method. (A2): Single trial (#1) for source *X*1 (−36, 52, 2) was color coded by the type of inversion method. (B1): 100 trials for source*X2* (−58, −16, 2) were averaged and color coded by the type of inversion method. (B2): Single trial (#1) for source *X2* (−58, −16, 2) was color coded by the type of inversion method. Red: Ground Truth; Blue: Inversion without fMRI spatial priors; Green: Inversion with all spatial priors including valid and invalid; Cyan: Inversion only with valid fMRI spatial priors; Magenta: Inversion with invalid spatial priors; Yellow: Virtual sensor technique from Beamformer.

**Fig. S7.**
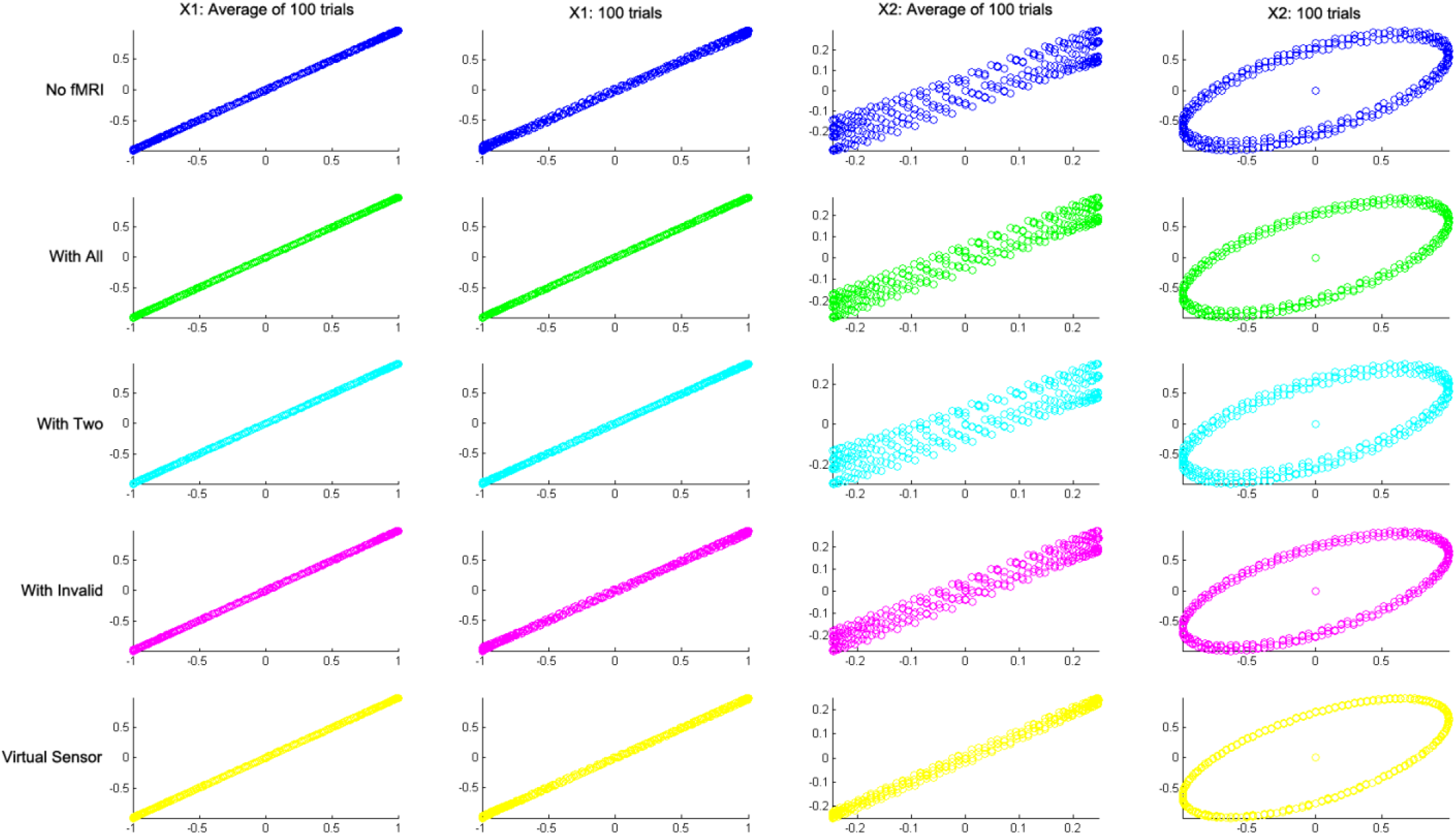
Scatter plots: the actual time courses for each time point and each trial were overlapped and plotted against the extracted time courses. The x axis represents the amplitude of the actual time courses that we constructed. The y axis represents the amplitude of the extracted time courses that we estimated. Blue: Inversion without fMRI spatial priors; Green: Inversion with all spatial priors including valid and invalid; Cyan: Inversion only with valid fMRI spatial priors; Magenta: Inversion with invalid spatial priors; Yellow: Virtual sensor technique from Beamformer.

**Fig. S8.**
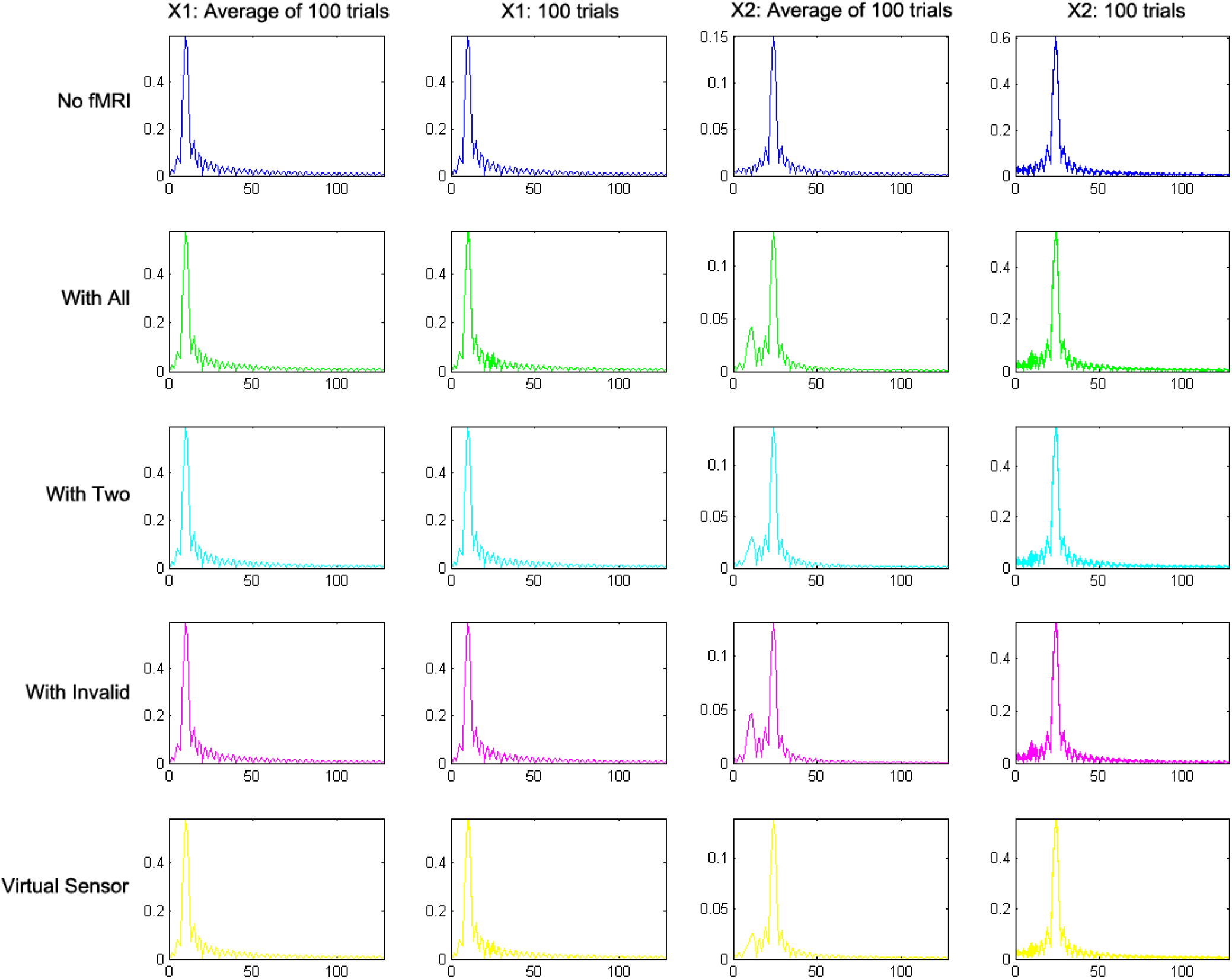
Single-sided amplitude spectrum plots: x axis is the frequency range from 0 to 128 Hz and y axis is the single-sided amplitude of the power spectrum. Blue: Inversion without fMRI spatial priors; Green: Inversion with all spatial priors including valid and invalid; Cyan: Inversion only with valid fMRI spatial priors; Magenta: Inversion with invalid spatial priors; Yellow: Virtual sensor technique from Beamformer.

## Notes

### Competing Interest Statement

The authors have declared no competing interest.

